# *LRRK2* G2019S mutation suppresses differentiation of Th9 and Treg cells via JAK/STAT3

**DOI:** 10.1101/2024.02.28.582561

**Authors:** Ningbo Zheng, Roshni Jaffery, Ashley Guerrero, Jiakai Hou, Yuchen Pan, Fang Zhou, Si Chen, Chunyu Xu, Nicholas A. Egan, Ritu Bohat, Ken Chen, Michael A. Schwarzschild, Xiqun Chen, Weiyi Peng

**Author notes:** Contributed equally (Ningbo Zheng, Roshni Jaffery). **Corresponding Authors:** Xiqun Chen, Massachusetts General Hospital, Harvard Medical School, Boston, MA.; Weiyi Peng, University of Houston, 3507 Cullen Boulevard, Houston, Texas 77204.

## Abstract

The Leucine-rich repeat kinase-2 (*LRRK2*) G2019S mutation, resulting in aberrantly enhanced kinase activity, is one of the well-recognized genetic risk factors in Parkinson’s Disease (PD). Increased LRRK2 activity was also observed in immune cells from PD patients. Emerging results have also unveiled an upsurge in α-synuclein (α-syn)-specific CD4^+^ T cell responses in PD patients. Given that *LRRK2* mutations in PD are germline mutations, there are unmet meets to explore whether *LRRK2* G2019S mutation contributes to the pathogenesis of PD via altering CD4^+^ T-cell functions. To fill this knowledge gap, we generated a new T cell receptor (TCR) transgenic mouse strain bearing *LRRK2* G2019S knock-in mutation, OT-II/LRRK2 (Refer to Mut). As CD4^+^ T cells from OT-II mice specifically recognize ovalbumin, this new strain enables us to explore the impact of *LRRK2* G2019S mutation on T-cell functions in an antigen-specific manner. We found that the abundance and proliferation of major immune subsets in spleen tissue from Mut mice are comparable to wild-type (OT-II, Refer to WT) control. However, when we characterized T cell differentiation in these two strains, T cells derived from Mut mice displayed increased Th2 differentiation (IL-4) and decreased Th9 (IL-9) and Treg (Foxp3^+^ %) differentiation. *LRRK2* G2019S mutation significantly altered the expression levels of master transcription factors (TFs) for T cell differentiation. Specifically, Mut T cells displayed an increase in mRNA expression of *Gata3* (TF for Th2), a decrease in expression of *Irf4* and *Foxp3* (TFs for Th9 and Treg, respectively). Mechanistically, *LRRK2* mutation decreased IL-9 production and Treg cell population through the JAK/STAT3 signaling. In conclusion, LRRK2 plays a critical role in regulating T cell differentiation, warranting further studies to evaluate the impacts of altered T cell differentiation led by *LRRK2* mutation in dopaminergic neuron damages.

## Introduction

Parkinson’s disease (PD) is a progressive neurodegenerative disease that affects both peripheral organs and the central nervous system (CNS), with neuroinflammation playing a critical role in its pathophysiology [1]. Although the PD community has begun to pay more attention to studying the immune system’s involvement, it has been primarily limited to evaluating the innate immune system, such as inflammatory cytokines and microglia, for decades [2]. These efforts lead to recognizing inflammation as a core feature of PD [3]. Recently, the presence of T cell infiltration in post-mortem brain sections was reported in PD patients [4]. Additionally, elevated levels of CD4^+^ and CD8^+^ T cells were observed in the substantia nigra pars compacta of PD patients compared to control subjects [5]. These findings spurred research into understanding the role of T cells in PD over the past five years. Emerging results have unveiled an upsurge in α-synuclein (α-syn)-specific T cell responses, primarily CD4^+^ T cells, in PD patients [6]. Given that the misfolding and aggregation of α-syn in the neuronal cells has been implicated in patients with neurodegenerative disorders [7], these results suggest that neuron-derived antigens could potentially elicit T-cell-mediated adaptive immune responses [6]. However, dysfunctions of T cells in PD patients, mainly related to PD risk factors, are largely unexplored.

Leucine-rich repeat kinase 2 (*LRRK2*) mutations are the strongest PD risk factors [8,9]. Through its kinase activity, LRRK2 modulates multiple intracellular signaling pathways, such as the MAPK/ERK pathways [10]. Hyperactivation of LRRK2 in neuron cells has been reported to disrupt various cellular functions, including vesicle trafficking, cytoskeletal dynamics, and mitochondrial function, which potentially contribute to the degeneration of dopamine-producing neurons in the brain [10]. Besides neuron cells, LRRK2 is also widely expressed in a broad range of immune cells, and upregulation of LRRK2 has been implicated in immune cells stimulated by microbial pathogens [11,12,13,14]. Although LRRK2 expression in T cells is relatively lower than in B cells and monocytes, T cells from PD patients display upregulated LRRK2 expression [12]. Additionally, most *LRRK2* mutations, such as the G2019S mutation, identified in PD patients are germline alterations and have been reported to lead to aberrantly enhanced kinase activity [15]. Therefore, it is reasonable to speculate that these PD-associated *LRRK2* mutations can also impact T cell function. However, whether T cells bearing *LRRK2* mutations exhibit altered function and contribute to the pathogenesis of PD remains unknown.

To fill in the knowledge gap connecting the LRRK2 pathway with T cell biology in the PD setting, we aim to comprehensively characterize the phenotypic and molecular consequences of *LRRK2* G2019S mutation in T cells. As most T cell epitopes for α-syn identified in PD patients are restricted by MHC class II [16], we mainly focused on CD4^+^ T cells in this study. Given that T cells express highly diversified T cell receptors (TCRs), which could confound T cell biology controlled by LRRK2, precise comparisons of T cell function in relation to *LRRK2* mutation require synchronizing TCRs to one epitope. Here, we generated a TCR transgenic mouse strain bearing *LRRK2* G2019S knock-in mutation, OT-II/LRRK2 (Refer to Mut). As TCRs expressed by T cells from OT-II mice specifically recognize ovalbumin (OVA) dependent on MHC class II molecules [17], this new strain enables us to explore the impact of *LRRK2* G2019S mutation on T cell functions and differentiation in an antigen-specific manner. Our immune profiling analysis of peripheral lymphoid tissues from Mut mice revealed that *LRRK2* mutation has limited impacts on the abundance of immune subsets. Proliferation of LRRK2 mutant-T cells is comparable with wild-type (OT-II, Refer to WT) T cells. However, we found that LRRK2 mutant-T cells exhibit a distinct preference for differentiating into different T helper (Th) cell types. Specifically, LRRK2 mutant-T cells displayed increased Th2 differentiation and decreased Th9 and Treg differentiation. Mechanistic studies further revealed that the JAK/STAT3 signaling mediates the role of LRRK2 in controlling the differentiation of CD4^+^ T cells. Our findings provide direct evidence to support T cell dysfunction in PD patients and shed new mechanistic insights on how *LRRK2* mutations are involved in PD initiation and progression.

## Materials and Methods

### Mice

By crossing OT-II (#004194) and *LRRK2* G2019S knock-in (#030961) mice obtained from The Jackson Laboratory (Bar Harbor, ME), we generated and bred OT-II mice bearing homozygous G2019S mutation of *LRRK2* (OT-II/LRRK2, Mut) mice in the specific pathogen-free barrier facility at the University of Houston (UH). OT-II (WT) mice were also maintained and used as controls in reported experiments. Mice were handled following protocols approved by the UH Institutional Animal Care and Use Committees. Both male and female mice were used.

### T cells differentiation

Naïve CD4^+^ T cells isolated from splenocytes by the EasySep Mouse CD4^+^ T-cell Isolation Kit (#19852, STEMCELL Technologies, Vancouver, Canada) were used to evaluate the capability to differentiate into different T cell subsets as previously described [18,19,20]. Purified naïve CD4^+^ T cells were stimulated with 2.5 μg/ml of anti-CD3e (#145-2C11, Cytek Biosciences, Fremont, CA) and 1 μg/ml of anti-CD28 (#553294, BD Biosciences, Franklin Lakes, NJ) antibodies in different culture conditions: (a) T0 condition (non-polarized): MEMα medium (#32-571-101, Fisher Scientific) supplemented with 10% FBS (#S11150, R&D Systems, Minneapolis, MN), 100U/ml of IL-2 (#NDC 65483-116-07, Prometheus Laboratories, San Diego, CA); (b) T1 condition (Th1-polarized): MEMα medium supplemented with 10% FBS, 100U/ml of IL-2, 4 ng/ml of IL-12 (#210-12, PeproTech, East Windsor, NJ), and 10 μg/ml of anti-IL-4 mAb (#BE0045, BioXCell, Lebanon, NH); (c) T2 condition (Th2-polarized): MEMα medium supplemented with 10% FBS, 100U/ml of IL-2, 10 ng/ml of IL-4 (#214-14, PeproTech), and 10 μg/ml of anti-IFN-γ mAb (#BE0055, BioXCell); (d) T9 condition (Th9-polarized): MEMα medium supplemented with 10% FBS, 100U/ml of IL-2, 10 ng/ml of IL-4, 1 ng/ml of TGF-β1 (#763104, BioLegend, San Diego, CA), 10 ng/ml of IL-1β (#401-ML/CF, R&D Systems), 10 μg/ml of anti-IFN-γ mAb. (e) TR condition (Treg-polarized): MEMα medium supplemented with 10% FBS, 300U/ml of IL-2, 5 ng/ml of TGF-β1, 10 μg/ml of anti-IFN-γ mAb, 10 μg/ml of anti-IL-4 mAb.

### Immune profiling

To characterize the immune profiles of peripheral lymphoid tissues from WT and Mut mice, single-cell suspensions of spleen tissues were prepared and depleted of red blood cells using ACK lysing buffer (#A1049201, Fisher Scientific). Splenocytes were incubated with antibodies against a panel of surface markers at 4°C for 30 minutes. Cells were then fixed and permeabilized using either the Foxp3/transcription factor staining buffer set (#00-5523-00, Fisher Scientific) or the BD fixation/permeabilization solution kit (#554714, BD BioSciences) according to the manufacturer’s protocols and then incubated with a cocktail of antibodies against intracellular markers. The antibodies used for staining included anti-CD3-PE (#555275, BD Biosciences), anti-CD4-eFluor 450 (#48–0042–82, eBioscience, San Diego, CA), anti-CD8-PE/Cy7 (#60–0081, TONBO Biosciences, San Diego, CA), anti-CD25-PerCP (#65-0251, TONBO Biosciences), anti-Foxp3-PE (#12-5773-82, Fisher Scientific), anti-CD19-APC (#20-0193, TONBO Biosciences), anti-Ki67-FITC (#11-5698-82, Fisher Scientific), anti-LRRK2-APC (#Ab195023, Abcam, UK). An LSRFortessa X-20 cell analyzer (BD Biosciences) was used for acquisition.

### ELISA

To determine the levels of cytokines produced by differentiated T cells, T cells after 5-day culture were collected, seeded into a new 96-well plate (1×10^5^ cells in 250 μl MEMα medium supplemented with 10% FBS per well), and restimulated with 50 ng/ml PMA (#P1585, Sigma-Aldrich, St. Louis, MO) overnight. The supernatant was collected and used for ELISA analysis, including mouse IFNγ (#DY485, R&D Systems), mouse IL-4 (#DY404, R&D Systems), and mouse IL-9 (#DY409, R&D Systems).

### mRNA expression analysis

Total RNA was extracted from T cells using the RNeasy Mini kit (#74104, Qiagen, Hilden, Germany) according to the manufacturer’s instructions. cDNA was synthesized using iScript Reverse Transcription Supermix (#1708841, Bio-Rad Laboratories, Hercules, CA). To determine mRNA levels of genes-of-interest (GOI), quantitative real-time polymerase chain reaction (qRT-PCR) using gene-specific primers and universal SYBR Green Supermix (#1725274, Bio-Rad Laboratories) was performed. Expression of each gene-of-interest was normalized to the expression of β-Actin (*Actb)*, a housekeeping gene, and triplication was performed for each sample. RNA-Seq libraries were prepared and subjected to paired-end sequencing by the Next-Generation Sequencing Facility at Novogene to characterize transcriptomic profiles of cultured T cells.

The primers used for qRT-PCR include m*Actb*-F: 5’-GGCTGTATTCCCCTCCATCG; m*Actb*-R: 5’-CCAGTTGGTAACAATGCCATGT; m*Tbx21*-F: 5’-CCACCTGTTGTGGTCCAAGTTC; m*Tbx21*-R: 5’-CCACAAACATCCTGTAATGGCTTG; m*Gata3*-F: 5’-CCTCTGGAGGAGGAACGCTAAT; m*Gata3*-R: 5’-GTTTCGGGTCTGGATGCCTTCT; m*Irf4*-F: 5’-GAACGAGGAGAAGAGCGTCTTC; m*Irf4*-R: 5’-GTAGGAGGATCTGGCTTGTCGA; m*Foxp3*-F: 5’-CCTGGTTGTGAGAAGGTCTTCG; m*Foxp3*-R: 5’-TGCTCCAGAGACTGCACCACTT; m*Lrrk2*-F: 5’-ATCTCACCCTTCATGCTTTCTG; and m*Lrrk2*-R: 5’-TCTCAGGTCGATTGTCTAAGACT.

### RNA-Seq Data analysis

Trimmomatic [21] was used for quality assurance to exclude low-quality reads from RNA-seq and remove adapter sequences improperly inserted in the reads. Then, raw reads were aligned to mouse genome reference build mm10 [22] using HiSAT2 v2.2.0 [23] with the default parameters. The gene-level read counts were calculated using HTSeq v0.11.1 [24] based on mouse transcriptome annotation. For each comparison, genes with low expression, defined as having counts per million (cpm) less than 2, were filtered out. The expression values were log2-counts per million (log CPM) transformed with the voom tool from Limma v3.50.3 [25], which reduces the skew and stabilizes the variance, making data suitable for downstream linear modeling in differential gene expression analysis, which was also performed by Limma.

Gene set enrichment analysis (GSEA) was then performed to identify molecular pathways impacted by *LRRK2* mutation in T cells. For each comparison, the enrichment analysis was performed by the GSEA function provided by the R package clusterProfiler (version 4.8.3) [26]; it aims to evaluate the potential biological functions of two conditions in each comparison group. The gene list used here was ranked by −log10 (p-value) * signed log (fold change). It utilizes the Kyoto Encyclopedia of Genes and Genomes (KEGG) pathway [27], Gene Ontology (GO) [28], and Hallmark gene sets in Molecular Signature Database (MSigDB) [29] to explore the enrichment from different aspects. The genome annotation information was retrieved with R package org.Mm.eg.db (version 3.8.2). In the GSEA method, the default permutation size is 1000. Gene ratio is the count of core enrichment genes divided by the count of pathway genes. A p-value was also derived from the permutation. Benjamini-Hochberg method was used to obtain adjusted p-values.

### Protein expression analysis

Western blot was performed to determine protein expression levels. An equal number of cells were collected and lysed in Laemmli Sample Buffer (#1610737, Bio-Rad) containing Pierce Protease and Phosphatase Inhibitor (#A32959, Fisher Scientific). 10-30 μl of protein lysates were used for the western blot analysis. Pierce Detergent Compatible Bradford Assay Kit (#PI23246, Fisher Scientific) was used to determine protein concentration. Antibodies used for western blot analyses include anti-LRRK2 (1:500, #13046, Cell Signaling Technology), anti-pLRRK2 S935 (1:500, #ab133450, Abcam), anti-β-Actin (1:4000, #sc-47778 HRP, Santa Cruz Biotechnology), anti-pSTAT3 (Tyr705) (1:500, #9131, Cell Signaling Technology), anti-pSTAT3 (Ser727) (1:500, #9134, Cell Signaling Technology), anti-STAT3 (1:1000, #9139, Cell Signaling Technology), anti-rabbit IgG (1:5000, #7074, Cell Signaling Technology), anti-mouse IgG, (1:4000, #7076, Cell Signaling Technology).

### Statistical analyses

The summary statistics (e.g., mean, SEM) are reported. The differences in continuous measurements between the two groups were assessed using a two-sample t-test posterior to data transformation (typically logarithmic, if necessary). A p-value of less than 0.05 was considered significant. Data acquisition and analysis from the cell analyzer were executed on FACSDiva and FlowJo softwares (BD Biosciences). Graph generation and statistical analyses were performed using GraphPad Prism software.

## Results

### Minimal changes in the immune profiles of peripheral lymphoid tissues were observed in OT-II/LRRK2 Mut mice

As resting T cells display lower LRRK2 expression compared with monocytes and B cells, the roles of LRRK2 in regulating T cells have not originally caught much attention. However, *LRRK2* mutations are well-recognized germline alterations associated with PD, and increased LRRK2 expression in T cells from PD patients was observed [12]. Furthermore, when we stimulated naïve CD4^+^ T cells isolated from OT-II (WT) mice by OVA peptide, we observed a dynamic expression of LRRK2 and phosphorated LRRK2 (p-LRRK2) at the protein level, with a peak at four days, in T cells in response to TCR stimulation (**Fig. 1A**). The findings from our group and others highlight the potential that *LRRK2* alterations in PD patients, such as *LRRK2* G2019S mutation, can result in T cell dysfunction. To test this possibility, we generated a new TCR transgenic mouse strain, OT-II/LRRK2 (Mut), by crossing OT-II and *LRRK2* G2019S knock-in mice. Age and gender-matched mice from the OT-II (WT) strain were included as controls. Notably, there were no significant differences in LRRK2 expression at the mRNA and protein levels in splenocytes between the WT and Mut mice (**Fig. 1B-C**). When we evaluated the abundance and proliferation of immune cells in the WT and Mut mice, we didn’t observe any significant changes in percentages of major immune subsets including CD4^+^, CD8^+^ T cells, and B cells (**Fig. 1D**). Additionally, as indicated by Ki67, a proliferation marker, Mut CD4^+^, CD8^+^ T cells, and B cells exhibit similar proliferative status as WT mice (**Fig. 1E**). These results suggest that the *LRRK2* G2019S mutation has minimal effects in T cell development and maintenance before antigen encounter.

**Fig 1.**
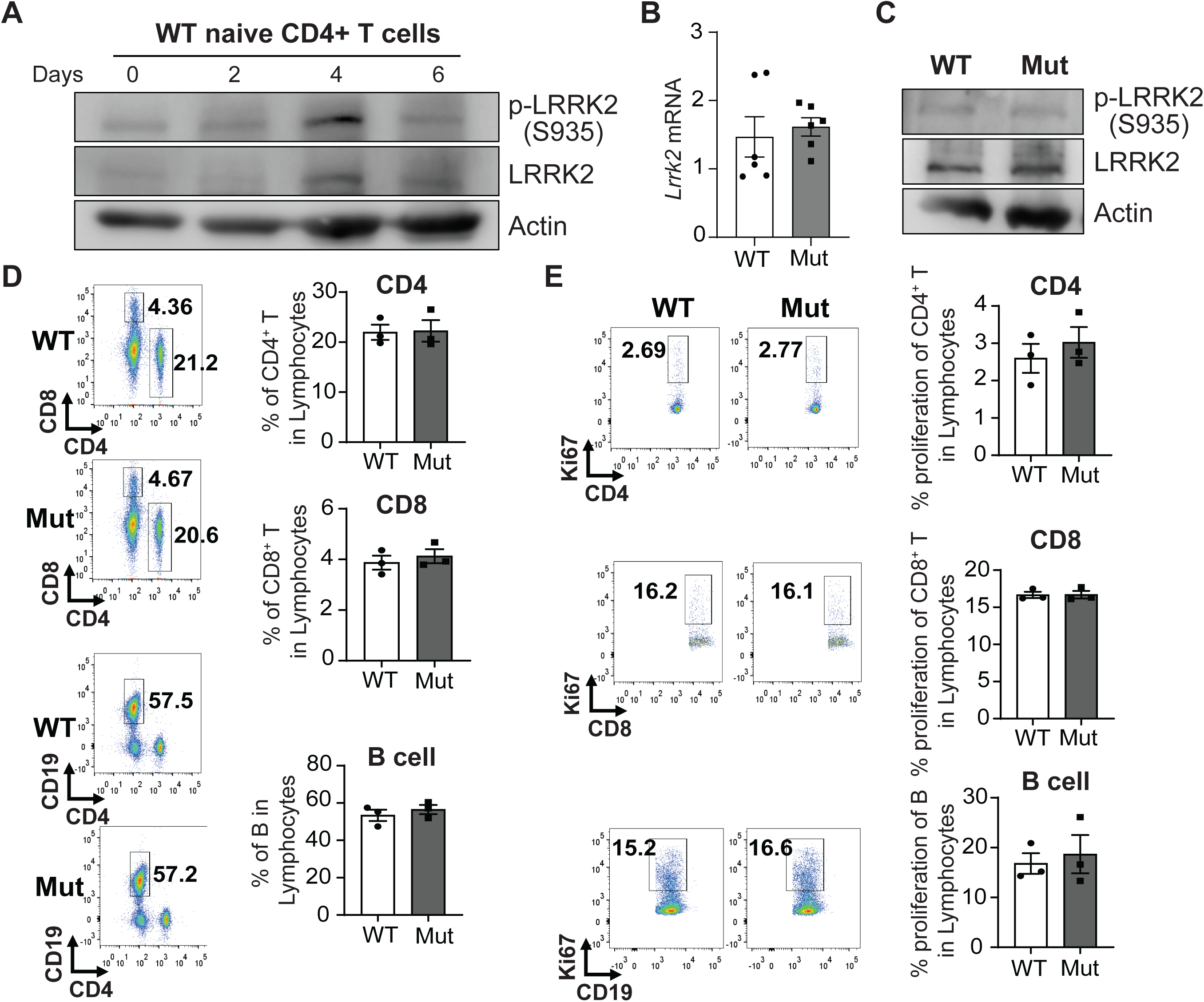
Splenocytes from OT-II/LRRK2 mutant mice exhibit minimal phenotypic changes. (**A**) Naïve CD4^+^ T cells were isolated from WT mice and stimulated with anti-CD3 Ab. Protein samples were collected from stimulated T cells at various time points (0-6 days). Changes of expression of LRRK2 and phosphorated LRRK2 (p-LRRK2) at the protein level in T cells during TCR stimulation. (**B**-**E**) Fresh splenocytes were isolated from WT and Mut mice. (**B**-**C**) *Lrrk2* mRNA expression levels in splenocytes were analyzed by RT-PCR (**B**), and protein levels were analyzed by Western blot (**C**). (**D**-**E**) Profiling of T and B cells was detected by flow cytometry analyses. Percentages of T cells (CD4^+^ and CD8^+^ T), B cells (CD4^−^ CD19^+^) (**D**), Ki67^+^ of T cells (CD4^+^ and CD8^+^ T), and B cells (**E**) were analyzed by flow cytometry analyses. Representative images and statistical results were shown. Data are mean ± SEM. Representative data from 3 independent experiments are shown. Triplications were included for all *in vitro* experiments.

### Mut T cells displayed distinct preferences for T cell differentiation

To further determine whether the *LRRK2* G2019S mutation affects CD4^+^ T cell differentiation, we isolated naïve CD4^+^ T cells from WT and Mut mice. These cells were then activated and cultured in various mediums to induce differentiation into Th1 (T1), Th2 (T2), Th9 (T9), and Treg (TR) cells. The Th0 (T0) culture medium containing only IL-2 was employed to sustain T cell growth without inducing polarization. Murine CD4^+^ T cells expanded in different types of polarization medium displayed variable LRRK2 expression. T cells under the T9 medium, which is required for the differentiation of IL-9-producing T cells, exhibit the highest LRRK2 expression among T cells differentiated in the rest types of polarization conditions (**Fig. 2A**). By evaluating the cell number changes of WT and Mut T cells under tested culture conditions, we found that the *LRRK2* G2019S mutation had no significant effects on T cell expansion in T cell polarization medium (**Fig. 2B**).

**Fig 2.**
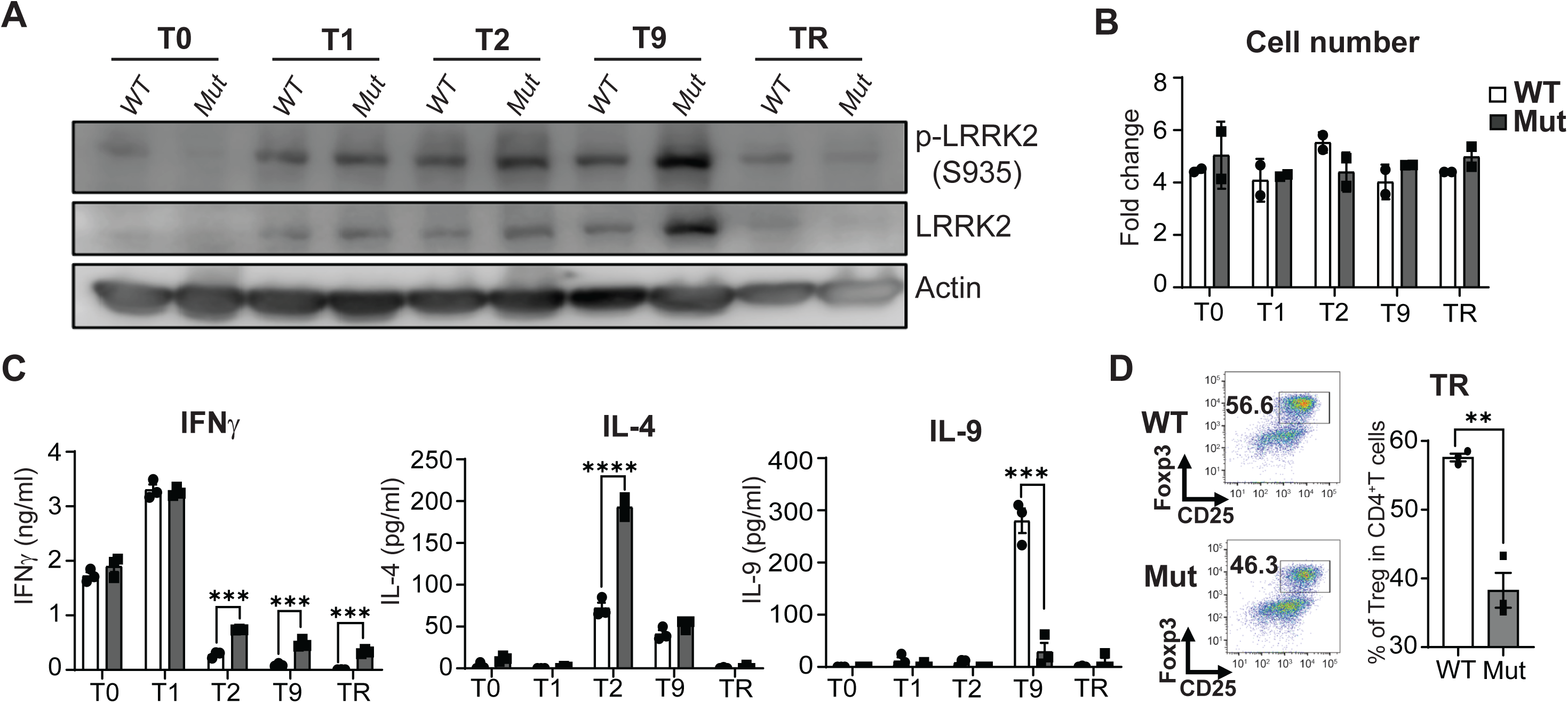
Mut T cells displayed distinct preferences for T cell differentiation. Naïve CD4^+^ T cells were stimulated with anti-mouse CD3/28 Abs under a set of T cell polarization conditions for Th1 (T1), Th2 (T2), Th9 (T9), and Treg (TR) differentiation for five days. The Th0 (T0) culture medium containing only IL-2 was employed to sustain T cell growth without inducing polarization. (**A**) Expression of LRRK2 and phosphorated LRRK2 (p-LRRK2) in T cells differentiated in different T cell polarization conditions were determined by Western blot. Representative figures from at least two independent experiments were shown. (**B**) The total cell number of different polarized CD4^+^ T cells under T0, T1, T2, T9, and TR conditions was counted by excluding trypan blue-positive dead cells. After differentiation for five days, (**C**) equal numbers of differentiated cells were seeded into a new 96-well plate (1×10^5^ cell/well) and restimulated with 50 ng/mL of PMA overnight. The levels of IFNγ, IL-4, and IL-9 in different condition mediums from overnight cultures were determined by ELISA, (**D**) the percentage of CD25^+^ Foxp3^+^ T cells in CD4^+^ T cells with or without *LRRK2* mutation differentiated under TR condition was determined by flow cytometry analysis. Representative flow cytometry results from three pairs of mice were shown. Data mean ± SEM. \*\**P*<0.01, \*\*\**P*<0.001, \*\*\*\**P*<0.0001. Triplications were included for all *in vitro* experiments.

Furthermore, IFNγ, IL-4, and IL-9, which are crucial effector cytokines for Th1, Th2, and Th9 cells, respectively, were selected as readouts to characterize the function of differentiated T cells. When we examined the cytokine levels produced by differentiated Mut and WT T cells, we observed significantly different cytokine levels in Mut T cells compared with WT T cells. Specifically, the IFNγ levels by Mut T cells under the T1 polarization condition are comparable with those by WT T cells (**Fig. 2C**). Increased IFNγ production by Mut T cells under T2, T9, and TR condition was observed (**Fig. 2C**). As expected, T2 condition induced the highest IL-4 production, and T9 condition results in the highest IL-9 production (**Fig. 2C**). More interestingly, Mut T cells under the T2 polarization condition exhibited an increased IL-4 production. At the same time, IL-9 secretion was notably reduced in Mut T cells under the T9 polarization conditions (**Fig. 2C**). Additionally, we also evaluated the percentage of Treg (Foxp3^+^ CD25^+^) in differentiated T cells under TR condition and found reduced Treg differentiation in Mut mice (**Fig. 2D**). Taken together, these results suggest LRRK2 plays a critical role in regulating T cell differentiation.

### Molecular changes in differentiated Th cells associated with *LRRK2* G2019S mutation

Given that Mut T cells exhibit distinct patterns in T cell differentiation, we analyzed the expression of master transcription factors (TF) for Th1 (*Tbx21)*, Th2 (*Gata3)*, Th9 (*Irf4)*, and Treg (*Foxp3)* in differentiated WT and Mut T cells. As shown in **Fig. 3A**, *LRRK2* G2019S mutation reduced the expression of *Irf4* and *Foxp3* in T cells differentiated in T9 and TR conditions, respectively, and increases *Gata3* expression in T cells differentiated in T2 condition (**Fig. 3A**). However, there is no significant change observed in the expression of *Tbx21* between WT and Mut T cells differentiated in T1 condition (**Fig. 3A**). These results are consistent with the observation that Mut T cells have reduced capability for Th9 and Treg differentiation, but increased Th2 differentiation.

**Fig. 3.**
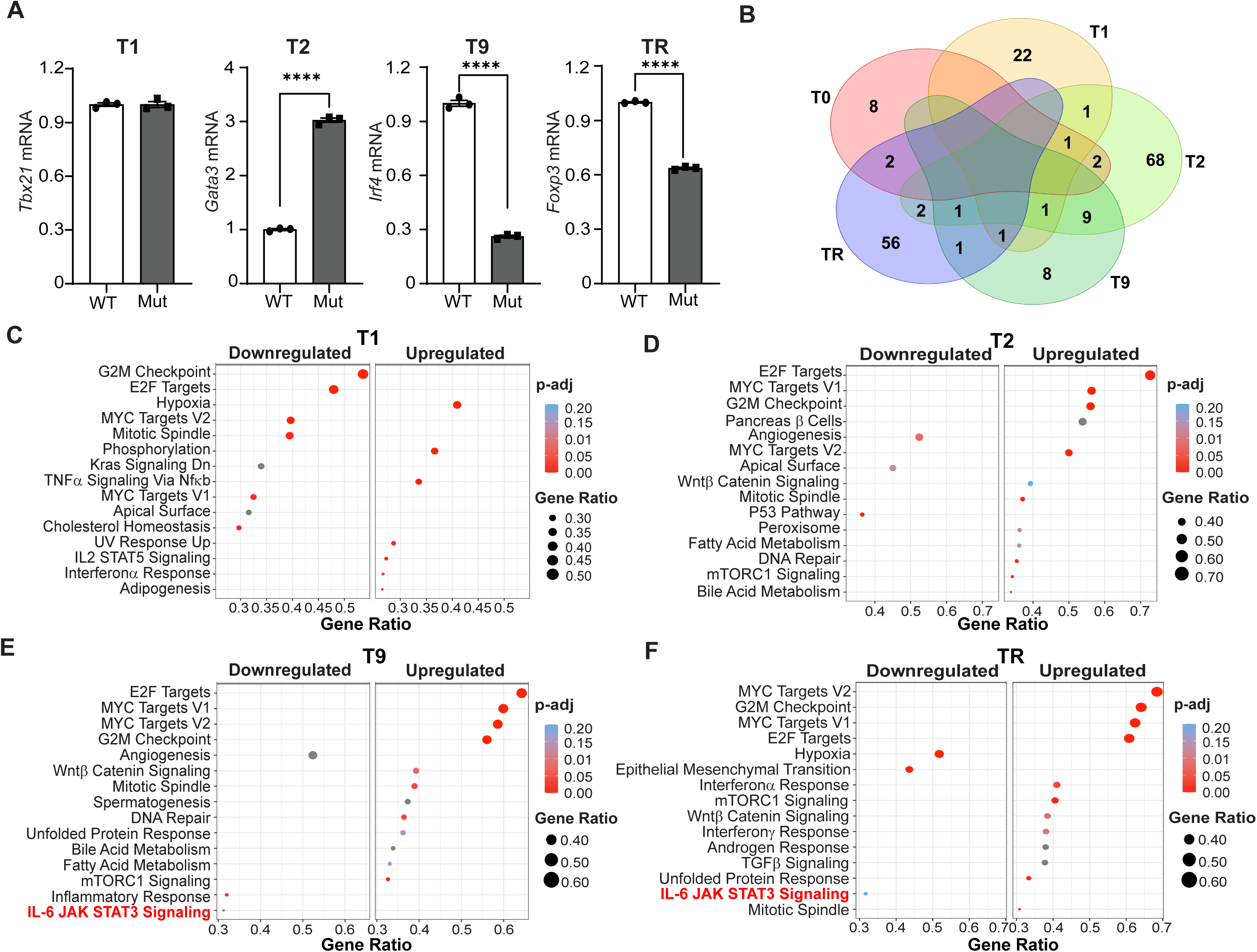
Molecular changes in differentiated Th cells associated with *LRRK2* G2019S mutation. Total RNA was extracted from WT and Mut polarized CD4^+^ T cells under T0, T1, T2, T9, and TR polarized conditions. (**A)** *Tbx21*, *Gata3*, *Irf4*, and *Foxp3* mRNA expression levels in T1, T2, T9, and TR polarized cells were detected by RT-PCR. (**B**) The Venn diagram shows the relationship between the WT and Mut T cells under T0, T1, T2, T9, and TR conditions. The number indicates the number of DEGs in each region. (**C-F**) Dot plots of GSEA results illustrating the Hallmark pathway enrichment of CD4^+^ T polarized cells under T1, T2, T9, and TR conditions from WT and Mut mice. Triplications were included for RT-PCR. Data mean ± SEM. \*\*\*\**P*<0.0001. Triplications were included for all *in vitro* experiments.

To further explore the mechanisms by which *LRRK2* G2019S mutation alters T cell differentiation, we performed RNA sequencing (RNA-seq) analyses to characterize the transcriptomic profiles of WT and Mut T cells differentiated under T1, T2, T9, and TR conditions. Significantly reduced *Irf4* expression in Mut T cells was also observed in RNA-seq results (**Supplementary Fig. 1**). By using the cutoff of adjust p-value (padj)<0.05 and |Fold change|≥1.25, differentially expressed genes (DEGs) in Mut T cells were identified. As shown in

**Fig. 3B**, the DEGs associated with the *LRRK2* mutation among T cells differentiated in different conditions are largely non-overlapped, with the greatest number of DEGs observed in T cells in T2 conditions. However, when we conducted the Gene Set Enrichment Analysis (GSEA) comparing the transcriptomic profiles between WT and Mut T cells, we found several enriched pathways across all tested four types of T cells, including MYC targets, E2F targets, and G2M checkpoints (**Fig. 3C-F**). It suggests that the *LRRK2* G2019S mutation can induce similar molecular changes in these pathways on different T subsets, although this role might be through distinct genes. More interestingly, the IL6-JAK-STAT3 signaling pathway was identified as one of the enriched differentially expressed pathways in T cells under T9 and TR conditions, not in T1 and T2 conditions (**Fig. 3C-F**). It implies that the JAK/STAT3 pathway might mediate the role of LRRK in T cell differentiation.

### LRRK2 modulates Th9 and Treg differentiation through the JAK/STAT3 signaling

To further explore whether the inhibition of IL-9 production and Treg differentiation by the *LRRK2* G2019S mutation through the JAK/STAT3 pathway, we evaluated the levels of STAT3 and phosphorylated STAT3 (p-STAT3) in T cells differentiated under T1, T2, T9, and TR conditions. The total STAT3 levels among differentiated T cells from Mut and WT mice are comparable cells (**Fig. 4A**). However, the levels of p-STAT3, the activated form, are variable among tested T cells, and when compared differentiated WT T cells, T cells under T9 condition display the lowest p-STAT3 (**Fig. 4A**). Interestingly, under T9 condition, the p-STAT3 levels in differentiated Mut T cells are significantly higher than WT T cells (**Fig. 4A**). Next, we used Stattic, a STAT3-selective inhibitor, to evaluate whether STAT3 inhibitor can rescue the phenotypic changes in T cell differentiation caused by the *LRRK2* G2019S mutation. Our results demonstrated that Stattic treatment can restore IL-9 level and *Irf4* mRNA level in Mut T cells differentiated under T9 condition (**Fig. 4B-C**). Moreover, Stattic treatment has no effect on the expression of *Foxp3* in WT T cells differentiated in TR condition, but upregulates *Foxp3* in Mut T cells differentiated in TR condition (**Fig. 4D**). These results demonstrate STAT3 inhibition can result in a full rescue in Th9 differentiation and a partial rescue in Treg differentiation observed in Mut T cells. In summary, our results showed that enhanced LRRK2 activity in T cells can promote STAT3 activation and lead to reduced Th9 and Treg differentiation (**Fig. 4E**).

**Fig. 4.**
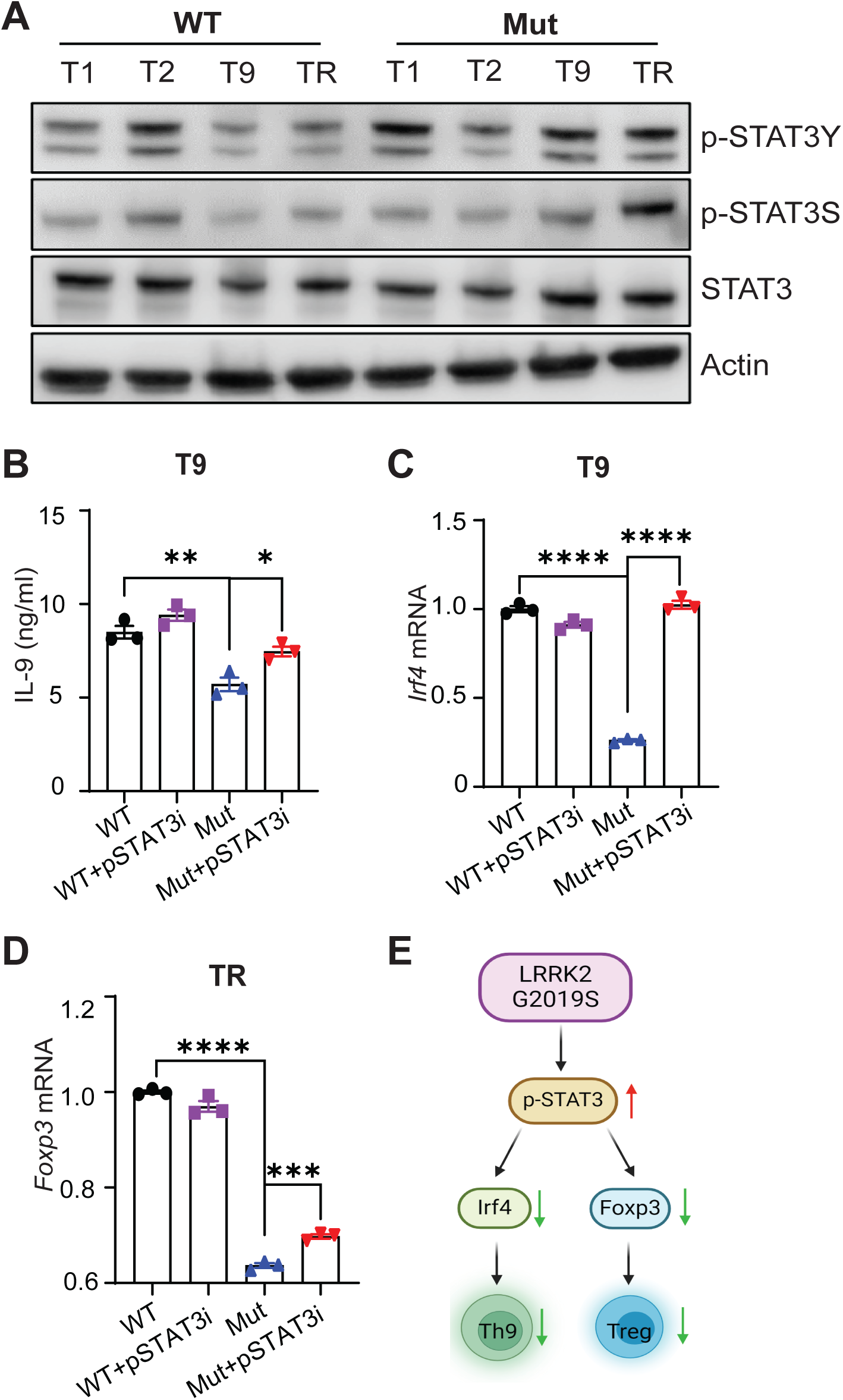
LRRK2 modulates Th9 and Treg differentiation through the JAK/STAT3 signaling. Naïve CD4^+^ T cells were purified from spleens of WT and Mut mice and cultured for five days in the presence of Anti-Mouse CD3e/CD28 antibodies with T1, T2, T9, and TR polarized cell conditions as previously described. (**A**) Total STAT3 protein and phosphorylation levels (p-STAT3Y, Tyr705 and p-STAT3S, Ser727) from WT and Mut polarized CD4^+^ T cells under T1, T2, T9, and TR polarized conditions were determined by Western blot. (**B**-**D**) Stattic, a STAT3 inhibitor. As described above, murine CD4^+^ T cells from WT and Mut mice were differentiated in T9 and TR conditions. For the groups receiving Stattic treatment, 1μM of Stattic was included in the culture medium after culturing 48 hrs. After a total of five days, (**B**) equal numbers of differentiated cells were seeded into a 96-well plate (1×10^5^ cells/well) and restimulated with 50 ng/mL of PMA overnight. The levels of IL-9 in culture medium from T cells under the T9 condition were measured by ELISA. (**C**) The *Irf4* mRNA level from T cells under the T9 condition were measured by RT-PCR. (**D**) The *Foxp3* mRNA level from the TR condition was measured by RT-PCR. (**E**) Schematic representation of signaling events depicting the impact of *LRRK2* G2019S mutation on downregulating Th9 and Treg cell differentiation. Enhanced LRRK2 activity associated with the G2019S mutation promoted STAT3 activation and led to reduced Th9 and Treg differentiation. Triplications were included for all *in vitro* experiments. \**P*<0.05 \*\**P*<0.01, \*\*\**P*<0.001 \*\*\*\**P*<0.0001.

## Discussion

Due to the presence of the blood-brain barrier (BBB), the central neuron system (CNS) has been recognized as one of the immune-privileged sites for a long time, particularly in the adaptive immune system. However, recent clinical trials showed encouraging results of T cell-mediated immunotherapy in treating patients with primary brain tumors. More importantly, both preclinical and clinical studies demonstrate a positive correlation between α-syn-specific T responses and the development of neurodegeneration diseases, particularly PD [6,16,30,31]. These new findings demonstrate that T cells can reach CNS and underscore the involvement of T cell immunity in dopaminergic neuron damage, which is currently underappreciated in the PD community. Here, we approached this question from a unique angle, namely, T cell dysfunction associated with genetic alterations in PD. By using a newly established TCR transgenic murine strain bearing the *LRRK2* G2019S mutation, we sought to reveal LRRK2-controlled cellular events in T cells. Although we found the *LRRK2* G2019S mutation has limited impacts on T cell proliferation, the *LRRK2* G2019S mutation significantly hindered Th9 and Treg differentiation, implying altered T cell responses in PD patients, especially in PD patients with *LRRK2* mutation. Distinct subsets of CD4^+^ T cells, such as Th1, Th2, Th9, and Treg cells, have been implicated to play eminently diverse roles in various human diseases. Th1 cells are critical in promoting cell-mediated immunity by producing cytokines like IFNγ, TNF-α, and IL-2. Through these effects, Th1 cells assist in the elimination of infected or aberrant cells to prevent infection and cancer. However, hyperactivated Th1 responses may lead to immune responses targeting self-antigens presented by healthy cells, resulting in autoimmune diseases [32]. On the other hand, Th2 cells, through the production of IL-4, IL-5, and IL-13, can trigger the activation of IgE-producing B cells, eosinophils, and mast cells and lead to type I hypersensitivity reaction [33]. Furthermore, tumor-specific Th2 cells were also reported to eradicate tumors by triggering an in situ inflammatory immune response [34]. Additionally, the interplay between Th1 and Th2 responses is crucial for maintaining immune balance and preventing pathological conditions [35]. IL-9, mainly produced by Th9 cells, can escalate immune responses by stimulating the production of other pro-inflammatory cytokines (such as IL-4, IL-5 and IL-13), and promote the activity of mast cells, CD8^+^ T cells, and natural killer (NK) cells. Moreover, Th9 cells can directly induce apoptosis in cancer cells and modulate the tumor microenvironment [36,37,38]. Thus, Th9 cells were initially recognized as an immune subset of intensified inflammatory and antitumor immune responses. However, Th9 cells also produce a large amount of IL-10, an immunosuppressive cytokine [39]. The pro-tumor roles of Th9 cells were also reported in recent studies. A correlation between Th9 tumor accumulation and poor survival was observed in lung cancer patients [40]. These results suggest a dual effect of Th9 cells in controlling immune responses. Unlike the above-mentioned subset, Tregs serve as a brake to suppress immune responses and maintain immune tolerance. Tregs produce immunosuppressive cytokines such as IL-10 and TGF-β [41]. The immunosuppressive receptors/ligand expressed on Treg can also impair cytotoxic T cells and NK cells. The presence of Treg in the tumor microenvironment is one of the tumor immune evasion mechanisms [42]. However, a noticeable gap exists in the current literature concerning their roles in the pathogenesis of PD. Recently, a study showed that reduced percentages of Treg and impaired Treg function correlate with PD severity [43]. Reduced serum IL-9 levels were observed in PD patients. [44,45,46]. Our results showed that *LRRK2* G2019S mutation can shift the balance of T cell differentiation among different subsets. Mut T cells display T cell dysfunctions associated with reduced preference for Treg and Th9 differentiation, consistent with clinical observations. Together with findings supporting the indispensable roles of Tregs and Th9 cells in autoimmune diseases, our results warrant future studies to determine whether LRRK2-associated T cell dysfunction modulates the magnitude of T cell-mediated neuron damages in the PD setting.

Mechanistically, we found that the role of LRRK2 in regulating Th9 and Treg differentiation is mediated by STAT3, a transcription factor controlling various inflammatory responses [47]. Previously, *LRRK2* gain-of-function mutations have been reported to enhance the activation of STAT3 signaling pathways and contribute to tumorigenesis in colon cancer [48]. Additionally, knocking down LRRK2 expression in tumor cells reduces the phosphorylation of STAT3 and ultimately suppresses tumor cell growth [49]. However, LRRK2 activity has not been linked with the activation of STAT3 in T cells. Our data indicates that the association of LRRK2 and STAT3 in T cells is similar to that in tumor cells. Previously, STAT3 has been recognized as a key player in controlling T cell differentiation. STAT3-deficient mice exhibit impaired Th2 differentiation, suggesting that STAT3 signaling promotes Th2 phenotype [50]. In contrast, STAT3 represses Th9 and Treg differentiation. T cell-specific STAT3 knockout mice have an increased percentage of IL-9-producing T cells after stimulation [51]. The effects of the LRRK2-STAT3 axis on Th2 and Th9 differentiation reported in our study are consistent with these findings. In the presence of TCR stimulation and TGFβ, both silencing STAT3 expression and STAT3 inhibitors were reported to result in downregulating Foxp3 expression in CD4^+^ CD25^-^ T cells, suggesting STAT3 activation could promote Treg differentiation [52]. However, more recent studies showed that STAT3 signaling plays a reciprocal role in controlling the development of Th17 cells and Tregs. Treg-specific ablation of STAT3 promotes Th17 differentiation and leads to the development of fatal intestinal inflammation [53]. In the presence of TGFβ and IL-6, STAT3 inhibitor treatment leads to an increased Treg population and decreased Th17 cells [54]. Our results show that increased STAT3 activation by *LRRK2* mutation reduces the Treg population in the TR polarization condition. The consequence of STAT3 activation in controlling Treg development could depend on the immune microenvironment, such as cytokine concentrations.

Taken together, we demonstrated that PD-associated *LRRK2* mutation can disturb the balance of T cell subset differentiation. Specifically, *LRRK2* G2019S mutation promotes Th2 but suppresses Th9 and Treg populations. Although we only evaluated one type of *LRRK2* mutation that occurred in PD, opposite effects of LRRK2 inhibition as *LRRK2* G2019S mutation on T cell differentiation were observed. We expect other gain-of-function *LRRK2* mutations, such as the R1441C mutation in the Ras of complex GTPase (ROC) domain [55], could lead to similar dysfunction in T cell differentiation. However, future studies are required to evaluate the contribution of LRRK2-associated T cell dysfunction to the pathogenesis of PD. Recently, we successfully generated a new transgenic strain in which OVA is expressed explicitly on dopaminergic neurons. By transferring WT and Mut-T cells in this strain, we can determine whether OVA-specific T subset cells play protective or offensive roles in dopaminergic neurons. Additionally, it is worthwhile to apply a similar analysis to define the effect of *LRRK2* mutations on CD8^+^ T cell proliferation, survival, differentiation, and cytotoxicity function. In conclusion, our research highlights that the PD-related risk factor, *LRRK2* G2019S mutation, alters CD4^+^ T cell differentiation and sheds new light on T cell involvement in PD development.

## Grant Support and Acknowledgment

This work was supported in part by the following grants: Aligning Science Across Parkinson’s [ASAP-00312] through the Michael J. Fox Foundation for Parkinson’s Research (M.A. Schwarzschild, X.C., W.P.); The University of Houston GEAR research funds kindly provided to W.P. For the purpose of open access, the authors have applied a CC-BY public copyright license to the Author Accepted Manuscript (AAM) version arising from this submission.

## Authors’ Contributions

**Conception and Design:** X. Chen and W. Peng

**Acquisition of data (provided required animals, cells, etc.):** N. Zheng, R. Jaffery, A. M. Guerrero, J. Hou, F. Zhou, S. Chen, C. Xu, R. Bohat, N. A. Egan and W. Peng

**Analysis and interpretation of data (statistical analysis and bioinformatics analysis):** N. Zheng, R. Jaffery, Y. Pan, K. Chen, M. A. Schwarzschild, X. Chen, W. Peng

**Writing and/or revision of the manuscript:** N. Zheng, R. Jaffery, M. A. Schwarzschild, X. Chen and W. Peng

**Study supervision**: W. Peng

