## Supplementary data for "*LRRK2* G2019S mutation suppresses differentiation of Th9 and Treg cells via JAK/STAT3"

[#]Contributed equally

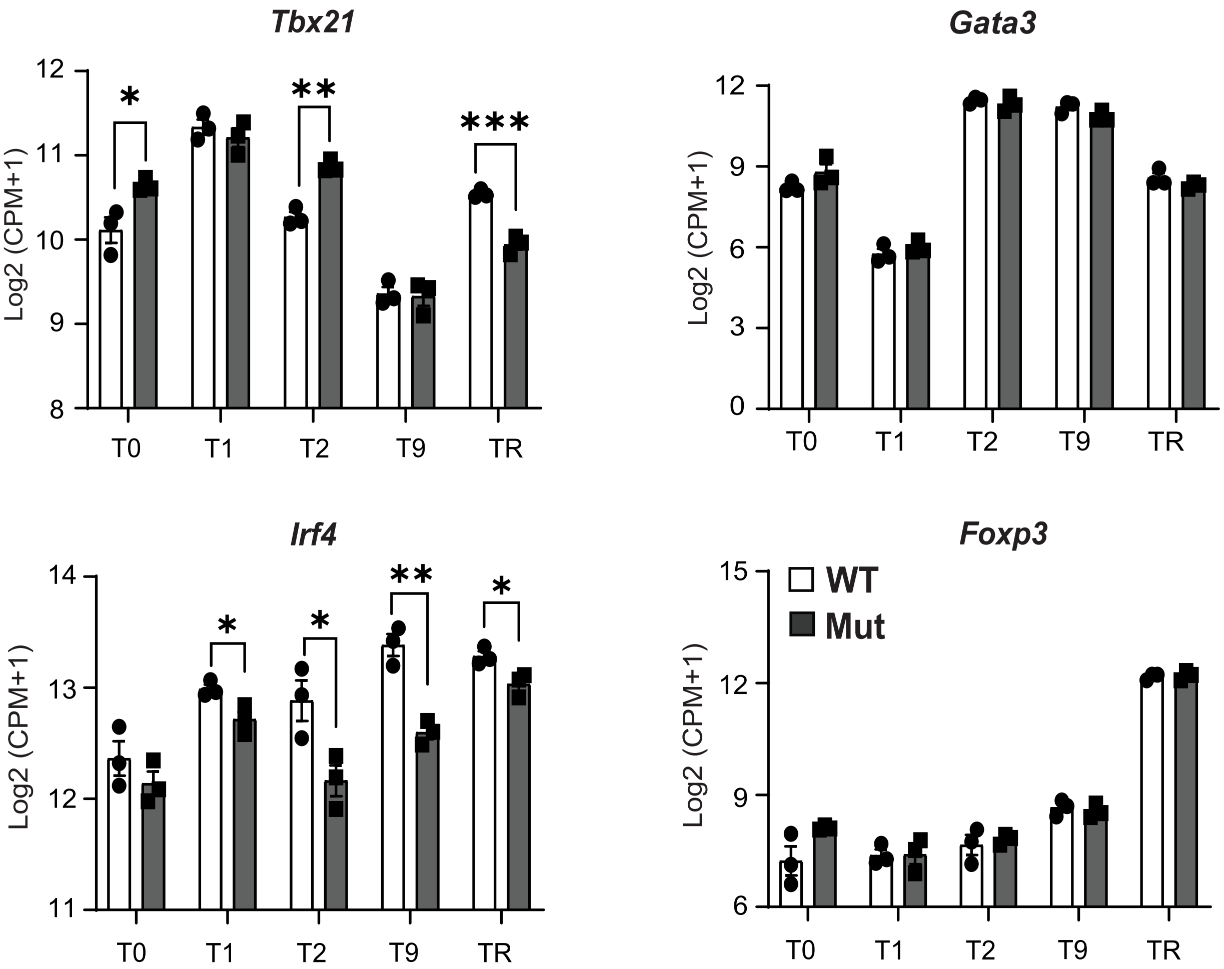

Tbx21
Log2 (CPM+1)
12
11
10
9
8
*
**
***
T0
T1
T2
T9
TR
Gata3
Log2 (CPM+1)
12
9
6
3
0
T0
T1
T2
T9
TR
Irf4
Log2 (CPM+1)
14
13
12
11
*
*
**
*
T0
T1
T2
T9
TR
Foxp3
WT
Mut
Log2 (CPM+1)
15
12
9
6
T0
T1
T2
T9
TR

### Figure legend

**Fig. S1. Molecular changes in differentiated Th cells associated with *LRRK2* G2019S mutation.** Total RNA was extracted from WT and Mut polarized $CD4^+$ T cells under T0, T1, T2, T9, and TR polarized conditions. *Tbx21*, *Gata3*, *Irf4*, and *Foxp3* mRNA expression levels in T0, T1, T2, T9, and TR polarized cells were detected by RNA-Seq. The logarithm base 2 transformation is applied to the CPM (counts per million) values. Data are mean ± SEM. *$P$<0.05, **$P$<0.01, ***$P$<0.001.
